# Comparative analysis of neutrophil and monocyte epigenomes

**DOI:** 10.1101/237784

**Authors:** Daniel Rico, Joost HA Martens, Kate Downes, Enrique Carrillo-de-Santa-Pau, Vera Pancaldi, Alessandra Breschi, David Richardson, Simon Heath, Sadia Saeed, Mattia Frontini, Lu Chen, Stephen Watt, Fabian Müller, Laura Clarke, Hindrik HD Kerstens, Steven P Wilder, Emilio Palumbo, Sarah Djebali, Emanuele Raineri, Angelika Merkel, Anna Esteve-Codina, Marc Sultan, Alena van Bommel, Marta Gut, Marie-Laure Yaspo, Miriam Rubio, José María Fernandez, Anthony Attwood, Victor de la Torre, Romina Royo, Stamatina Fragkogianni, Josep Lluis Gelpí, David Torrents, Valentina Iotchkova, Colin Logie, Ali Aghajanirefah, Abhishek A Singh, Eva M Janssen-Megens, Kim Berentsen, Wendy Erber, Augusto Rendon, Myrto Kostadima, Remco Loos, Martijn A van der Ent, Anita Kaan, Nilofar Sharifi, Dirk S Paul, Daniela C Ifrim, Jessica Quintin, Michael I. Love, David G Pisano, Frances Burden, Nicola Foad, Samantha Farrow, Daniel R Zerbino, Ian Dunham, Tacow Kuijpers, Hans Lehrach, Thomas Lengauer, Paul Bertone, Mihai G Netea, Martin Vingron, Stephan Beck, Paul Flicek, Ivo Gut, Willem H Ouwehand, Christoph Bock, Nicole Soranzo, Rodericw Guigo, Alfonso Valencia, Hendrik G Stunnenberg

## Abstract

Neutrophils and monocytes provide a first line of defense against infections as part of the innate immune system. Here we report the integrated analysis of transcriptomic and epigenetic landscapes for circulating monocytes and neutrophils with the aim to enable downstream interpretation and functional validation of key regulatory elements in health and disease. We collected RNA-seq data, ChIP-seq of six histone modifications and of DNA methylation by bisulfite sequencing at base pair resolution from up to 6 individuals per cell type. Chromatin segmentation analyses suggested that monocytes have a higher number of cell-specific enhancer regions (4-fold) compared to neutrophils. This highly plastic epigenome is likely indicative of the greater differentiation potential of monocytes into macrophages, dendritic cells and osteoclasts. In contrast, most of the neutrophil-specific features tend to be characterized by repressed chromatin, reflective of their status as terminally differentiated cells. Enhancers were the regions where most of differences in DNA methylation between cells were observed, with monocyte-specific enhancers being generally hypomethylated. Monocytes show a substantially higher gene expression levels than neutrophils, in line with epigenomic analysis revealing that gene more active elements in monocytes. Our analyses suggest that the overexpression of c-Myc in monocytes and its binding to monocyte-specific enhancers could be an important contributor to these differences. Altogether, our study provides a comprehensive epigenetic chart of chromatin states in primary human neutrophils and monocytes, thus providing a valuable resource for studying the regulation of the human innate immune system.

## INTRODUCTION

Human neutrophils and monocytes are the most abundant nucleated myeloid cells in peripheral blood and are essential elements of the innate immune system that provide the first line of defense against infection (Dale et al., 2008). Neutrophils represent approximately 50-60% of leukocytes in blood, while approximately 1-5% are monocytes. During hematopoiesis both neutrophils and monocytes are derived from the same progenitor, the myelomonocytic progenitor (CFU-GM). Despite this and the fact that both have similar functions as phagocytes in their clearance of microbial pathogens and cytotoxic activity, these cells differ in many of their specific biological functions.

Neutrophils are mature terminally differentiated cells with a very short survival time that are key in the innate immune response to acute inflammation. They are recruited to the site of infection and kill bacterial and fungal pathogens by phagocytosis and by releasing reactive oxygen species and antibacterial proteins in order to destroy pathogens in surrounding tissues that the neutrophils have infiltrated. Under certain conditions, they also release the so-called neutrophil extracellular traps (NETs) that originate from their chromatin (Kolaczkowska et al., 2013; Mantovani et al., 2011).

Monocytes have multiple roles in innate and adaptive immunity. They are also recruited into the extravascular tissues where they rapidly differentiate into macrophages or dendritic cells that then remain for weeks to months within the local tissue environment and fulfill highly specialized cellular functions (Galli et al., 2011). Monocytes themselves perform phagocytosis; on activation they elicit a proinflammatory cytokine response (Fairfax et al., 2014) and present antigen, thus may also initiate the adaptive immune response in T cells (Randolph et al., 2008).

Neutrophils and monocytes are involved in human diseases including bone marrow failure or chonic neutropenia syndromes, as well as myeloproliferative disorders or myeloid leukemias (Dale et al., 2008). Moreover, monocytes and derived cells have been implicated in multiple autoimmune diseases including systemic lupus erythematosus (Marshak-Rothstein et al., 2006) and multiple sclerosis, where monocytes may be key in the response to IFN-β treatment (Zula et al., 2011).

As both monocytes and neutrophils circulate in the blood, their epigenome is directly influenced by the presence of factors such as inflammatory agents, nutrients and metabolites. The rapid turnover of this class of blood cells makes them suitable targets for epigenetic drugs. Indeed, HDAC and SIRT inhibitors are suggested to have a marked effect on monocyte function (and possibly neutrophil function) including migration, autoimmune responses and attenuation of inflammatory responses (Orecchia et al., 2011; Adcock, 2007). However, to refine these epigenetic modulation strategies, a detailed definition of the epigenetic state of these cell types is essential (Ostuni et al, 2016).

Although the morphological and functional differences between these common immune cells have been extensively studied during the last century, our current knowledge on the molecular determinants that drive these different phenotypes is very limited. To understand how these different cells arise and to characterize their role in different pathological conditions, it is important to investigate differences in their gene expression programs in the broader context of their common and cell-specific epigenetic characteristics.

Previous studies have set out to investigate specific epigenetic features in monocytes and neutrophils (Ostuni et al, 2016). In neutrophils, DNA methylation has been studied using whole-genome bisulfite sequencing (WGBS) at a 10x resolution in a pool of 6 healthy individuals (Hodges et al., 2011) and by 450K microarrays Ronnerblad et al., 2014). In monocytes, subsets of histone modifications have been profiled (Schmidl et al., 2014; Pham et al., 2012; Pham et al., 2013) as well as the DNA methylome using enrichment procedures such as methylated DNA immunoprecipitation (MeDIP) (Salpea et al., 2012; Shen et al., 2013). More recently, the International Human Epigenome Consortium (IHEC) has generated released >7000 reference epigenomic datasets (Stunnenberg et al, 2016) including these cell types.

Here, as part of the BLUEPRINT consortium (Abbott, 2011; Adams et al., 2012; Martens and Stunnenberg, 2013; www.blueprint-epigenome.eu) we present a comparative analysis of IHEC reference epigenome maps for human adult and cord blood monocytes and neutrophils which are already available to the scientific community (Tables S1-4). The data produced by BLUEPRINT for peripheral blood and cord blood monocytes and neutrophils includes:

* ChIP-seq data for H3K4me1 (H3 lysine 4 monomethylation), H3K4me3 (H3 lysine 4 trimethylation), H3K9me3, H3K27me3, H3K36me3 and H3K27ac (H3 lysine 27 acetylation).
* High-resolution DNA methylation maps of the neutrophil and monocytes by whole genome bisulfite sequencing (WGBS) at >50x coverage
* Transcriptome characterization through strand specific RNA-seq on ribo-depleted RNA.

Together, these data represented the first complete reference epigenomes for these two cell types as defined by the International Human Epigenome Consortium (IHEC) (www.ihec-epigenomes.org).

Bioinformatic analysis of this comprehensive dataset (Table S5) revealed that any single epigenetic feature, be it DNA methylation, a histone modification or the transcriptional profile could distinguish monocytes from neutrophils. The combinatorial analysis of chromatin marks showed that monocytes generally have more promoters with active marks and have a higher number of non-promoter regulatory active regions (enhancers) compared to neutrophils. Neutrophils, on the contrary were found to have a larger proportions of heterochromatin as well as regions with no marks. Although the DNA methylomes of these cell types were overall very similar, most differences in DNA methylation were found to overlap enhancer regions, suggesting DNA methylation could be used to modulate their activity. Transcriptome profiling by RNA-seq showed higher transcriptional activity in monocytes compared to neutrophils, with more genes overexpressed in monocytes. These results suggest that monocytes, as precursor cells with diverse differentiation potential, have a more plastic and active epigenome than the terminally differentiated neutrophils.

## RESULTS

### Monocytes display more active promoters and enhancers than neutrophils

We generated genome-wide maps for histone modifications associated with diverse regulatory and epigenetic functions. These include histone H3 lysine 4 mono- and trimethylation (H3K4me1 and −me3), H3K9me3, H3K27me3, H3K36me3 and H3K27ac. We profiled these histone marks in neutrophils and monocytes from the peripheral blood of four adults (AB) and from the umbilical cord blood of two newborns (CB) (Fig. 1A-C, Fig. S1-2). Principal component analysis (PCA) analysis revealed that histone modification patterns within a given cell type cluster together. Similar results were obtained with DNA methylation or gene expression data (Fig. 1D). Given the similarities between peripheral AB and CB for monocytes and neutrophils we have focused our analyses on comparing peripheral adult blood monocytes and neutrophils.

**Fig. 1.**
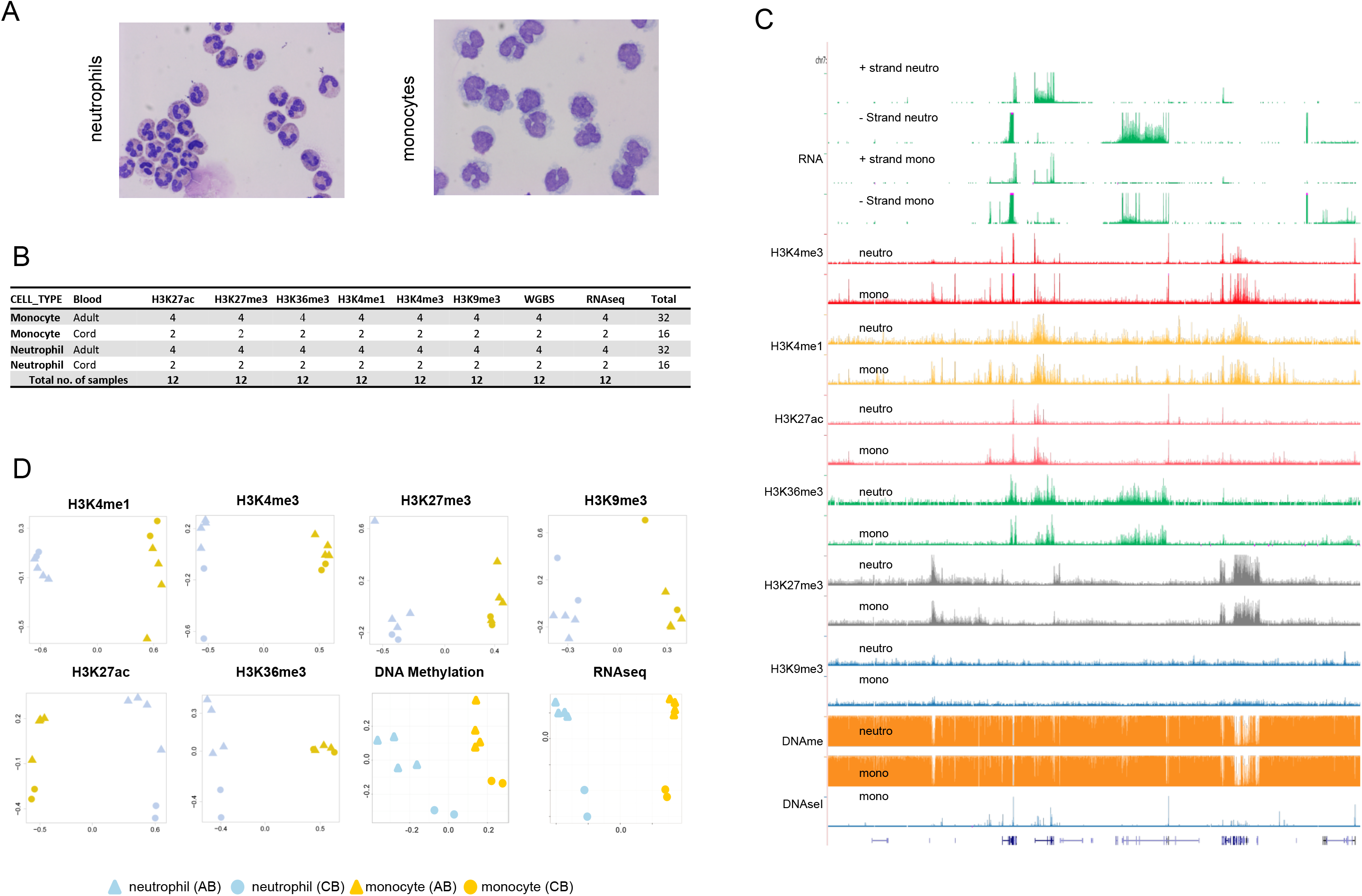
Overview of the data generated in this study. A. Cell images of neutrophils and monocytes isolated from peripheral blood. B. Summary table of samples assayed by each technology. In total 72 ChIP-seq tracks, 12 WGBS and 12 RNA-seq tracks were analyzed. C. Overview of a genomic region (chr7:25,447,050-27,726,032) showing representative RNA-seq, ChIP-seq and WGBS tracks in monocytes and neutrophils. In addition, a representaive track for DNAseI-seq using monocytes has been included. D. PCA analysis of histone modifications, DNA methylation and RNA-seq for cord and peripheral monocytes and neutrophils.

We used a Hidden Markov model approach (ChromHMM) for unsupervised segmentation of the genome into chromatin states (Ernst and Kellis, 2010). Several models searching for the optimal number of different states were generated, and a model with 11 different chromatin states was eventually selected as it offered the maximum number of states with biological interpretability (Fig. 2A, S4-5 and Table S6). Note that state 2 is characterized by the absence of marks tested here. As some of these states were functionally related, the 11 states were collapsed into five biological groups: **(1) Repressive heterochromatin (RHet)**, characterized by the enrichment of H3K9me3 (state 1), the enrichment of H3K27me3 (state 3) or the absence of any signal (state 2). RHet regions tend to be devoid of gene expression (~0.5 fold compared to the genome average, Table S6) and DNase I signal (~0.2 fold) in monocytes. (2) Active promoters (Apro) marked with H3K4me3, either alone (state 6), in combination with H3K4me1 (states 5) or with H3H27ac (state 7). Apro regions frequently overlap with a TSS (~48 fold enrichment). **(3) Repressed promoters (Rpro, state 4)** with H3K4me3, H3K4me1 and the H3K27me3 - also called *bivalent* or *poised* - that show lower accessibility by DNaseI (~9 fold) than Apro regions. (4) Regulatory elements (RegE) with H3K4me1 alone (state 9) or in combination with H3K27ac (state 8). RegE regions are enriched in DNase I (~22 fold) but show a lower enrichment in TSS (~2 fold) than in Apro (48 fold). These elements likely function as *latent* or *active* enhancers. **(5) Transcribed regions (TranR, states 10 and 11)** marked with H3K36me3 that show a clear enrichment in gene expression measured by RNA-seq (4 fold compared to the genome average).

**Fig. 2.**
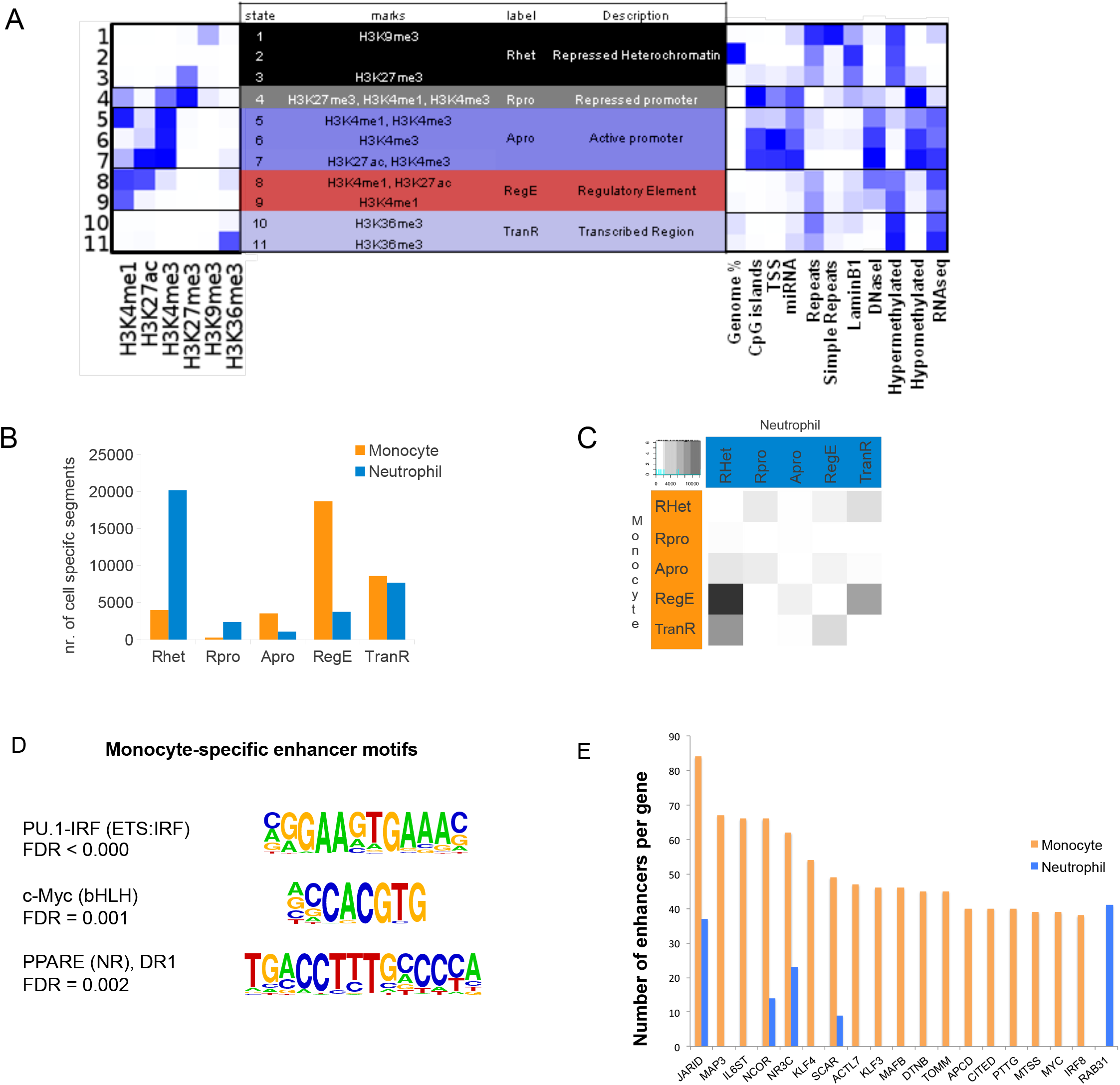
Chromatin state differences between neutrophils and monocytes. A. ChomHMM segmentation analysis using the 6 histone modification profiles in neutrophils and monocytes from peripheral blood. After testing different ChromHMM models, 11 different chromatin states were identified and named according to their presumed function. The heatmap on the left represents the relative probabilities of each histone mark in each state. The heatmap on the right shows the relative fold enrichment of overlap with each of the indicated genomic and epigenomic features. B. Number of genomic regions covered by cell-specific chromatin states in neutrophils and monocytes. C. Transitions between chromatin states in neutrophils and monocytes. Grey spots represent relative enrichment in the percentage of transitions. D. Representative motifs enriched in monocyte-specific regulatory regions. E. Enhancer scores for monocyte (orange) and neutrophils (light blue) for the top 20 genes with the highest absolute difference between enhancer scores in the two cell types. The enhancer score is defined as the number of regulatory regions associated to each gene by GREAT.

To first characterize cell-specific and shared putative regulatory elements, we compared the chromatin landscapes of adult monocytes and neutrophils by identifying regions that showed a consistent state in all four biological replicates for each cell type, while having a different state in the other cell type. We focused on the five functional groups described above (Fig 2B). Five times more repressive chromatin (RHet) segments were inferred in adult blood neutrophils compared to monocytes (Fig. 2B). In contrast, a greater number of regulatory elements (RegE, 5-fold) and active promoters (Apro, 4-fold) were inferred in monocytes. This difference in enhancer-like RegE segments matched the higher number of loci occupied by H3K27ac and H3K4me1 in monocytes observed in the differential analysis of single modifications between the two cell types (Fig. S6). Monocytes were enriched for transcribed regions (TranR) and regulatory elements (RegE), while the same genomic regions tended to encompass heterochromatic (RHet) regions in neutrophils (Fig. 2C). Interestingly, CB monocytes or neutrophils had a higher number of RegE regions than their AB counterparts (Fig. S7). Together, these results show that there is a marked difference in usage of specific epigenomic segments in neutrophils and monocytes. Neutrophils have far less marks associated with active transcription, as it can be expected for terminally differentiated cells. Based on their chromatin signature, monocytes can be considered lineage-committed multipotent cells, as they can further differentiate into macrophages, dendritic cells or osteoclasts.

We further sought to explore whether cell type-specific gene expression was associated to specific enhancer activity (Hnisz et al., 2013; Andersson et al., 2014). We partitioned chromatin segments labeled as enhancers (RegE) in monocytes and neutrophils into three groups: monocyte-specific (18,679), neutrophil-specific (3,749) and common enhancers (12,027). We found that enhancers were enriched both upstream and downstream annotated TSSs, with a prevalence in intronic regions (Fig. S8). We also searched for transcription factor binding sites (TFBS) sequences that were enriched in the neutrophil-specific and monocyte-specific enhancers. While no significantly enriched motifs were found in the neutrophil-specific enhancers, we identified many factors that potentially could bind to the monocyte-specific enhancers (False Discovery Rate [FDR] < 0.05, Table S7). Among them were TF motifs known to be important for monocyte and macrophage biology, such as PPARG, PU.1 and IRF (Fig. 2D) and interestingly also for pluripotency factors, including MYC.

It has been suggested that clusters of enhancers drive expression of genes that define cell identity (Hnisz et al., 2013). To identify such genes in monocytes and neutrophils, we calculated the number of putative enhancers for each gene and computed the difference between the two cell types. Of the top 20 genes with highest absolute difference, 19 showed more enhancers in monocytes, including NCOR2, IRF8 and MYC (Fig 2E, S10). Remarkably, not only MYC binding motifs were enriched in monocyte-specific enhancers (Fig. 2D), but also the MYC gene itself was associated with 39 enhancers in monocytes, while none of these enhancers were associated with MYC in neutrophils (Fig. 2E). Finally, we investigated functional clustering of genes corresponding to the inferred neutrophil-specific and monocyte-specific enhancers, using functional enrichment analysis implemented in GREAT (McLean et al., 2010). Neutrophil-specific enhancers were enriched in pathways related with p53, IL4, IL8, TNF and IFN signaling (Fig S9). Among the signaling pathways enriched in monocyte-specific enhancers were NFAT, IL2, IL12, CD40 and FAS.

Overall, these analyses suggest a greater plasticity of monocyte epigenomes compared to neutrophils. The data generated represents a large catalog of putative cell-specific and shared regulatory domains for further investigation, highlighting a host of known and novel pathways and transcriptional regulators for further study.

### Whole Genome DNA methylation maps and chromatin states

To establish comprehensive, high-resolution DNA methylation maps of the neutrophils and monocytes, we performed whole genome bisulfite sequencing with a median sequencing coverage of 63x per sample in a total of 12 samples (Fig. 1B). DNA methylation levels were inferred from the aligned reads using an algorithm that distinguishes between bisulfite-induced C-to-T conversion and genetic C-to-T differences (Kulis et al., 2012). Across all samples, we obtained DNA methylation estimates for an average of 23.68 million CpG sites. Details of the sequencing and calling statistics by sample are shown in Table S8. DNA methylation levels for a common set of 18.9 million CpG sites could be estimated in all samples, which forms the basis for the in-depth analysis reported below. In contrast, we did not observe any sites of consistent non-CpG methylation (Ziller et al., 2011) across replicates, and therefore our analyses focus exclusively on CpG methylation.

Consistent with previous observations in purified mouse hematopoietic cell populations (Ji et al., 2010; Bock et al., 2012), global DNA methylation levels were highly similar between neutrophils and monocytes (Fig. S11). Nevertheless, DNA methylation differences at the CpG level are highly informative, as PCA analysis of the DNA methylation profiles allowed the clear separation not only between neutrophils and monocytes, but also between cord blood and adult blood (Fig. 1D). In our segmentation analysis we showed that the DNA methylation status is very different in the chromatin state segments defined above (Fig. 2A). Our analysis revealed DNA hypermethylation at heterochromatin states 1 and 2 as well as at transcribed gene bodies (states 10 and 11), in line with previous reports (Lister et al., 2009, Brinkman et al, 2012). In contrast, hypomethylation was detected at active promoter segments, while regulatory elements showed intermediate to high levels of DNA methylation (Fig. 2A). To assess if the differences in DNA methylation at enhancer elements are linked to accessibility of the enhancers, we examined DNA methylation in enhancer regions overlapping with DHSs (available only for monocytes), compared to those that do not overlap. Interestingly, DNA methylation at accessible enhancer regions (identified with DNase I data) in monocytes was lower than at non-accessible regions (FDR < 10^−16^, T-test, Fig. S12), corroborating previous observations (Stadler et al., 2011) and suggesting that chromatin accessibility and DNA methylation can further refine subclasses of enhancers.

To further understand the relationship between DNA methylation and chromatin states, we tested differential methylation in DNA segments defined by the chromatin state patterns between samples. For this analysis, we used RnBeads software to test for differential methylation in each segment (Fig. S13) (Assenov et al., 2014). This approach has the advantage of increasing the statistical power in DMR detection, as it combines statistical evidence from neighboring CpGs in each DNA segment previously defined (Bock et al., 2012), as well as facilitating the biological interpretation of the significant DMRs. We compared neutrophils and monocytes using RnBeads and detected 17,129 DMRs (FDR < 0.01, beta difference > 0.1) comprising 4,62 Mb of the human genome, with approximately two thirds of the DMRs (11,429) showing higher methylation levels in monocytes. About half of the DMRs (53%) fall into introns, with only 34% of the DMRs located outside genes (Fig. S8).

Inspection of the DMRs in the context of chromatin states revealed that these are most enriched in regions that are enhancer/regulatory regions (RegE) in monocytes and neutrophils Fig. 3A-F) (FDR = 2 × 10^−46^, Chi-squared test), or enhancers in monocytes and heterochromatic (Rhet) in neutrophils (FDR = 2 × 10^−20^, Chi-squared test) (Fig. 3E-H). We observed 3.5 fold more hypermethylated DMR elements at regulatory regions (RegE) in monocytes compared to neutrophils, likely correlating with presence of these elements in introns of actively transcribed (and high in DNA methylation) genes (Lister et al., 2009). In contrast, the majority of RegE regions in monocytes that are marked as RHet in neutrophils, tend to be hypomethylated in monocytes (16 fold, Fig. 3E-G Fig S14). TSS methylation did not differ substantially between monocytes and neutrophils even when separating promoters in different configurations (Fig. S15).

**Fig. 3.**
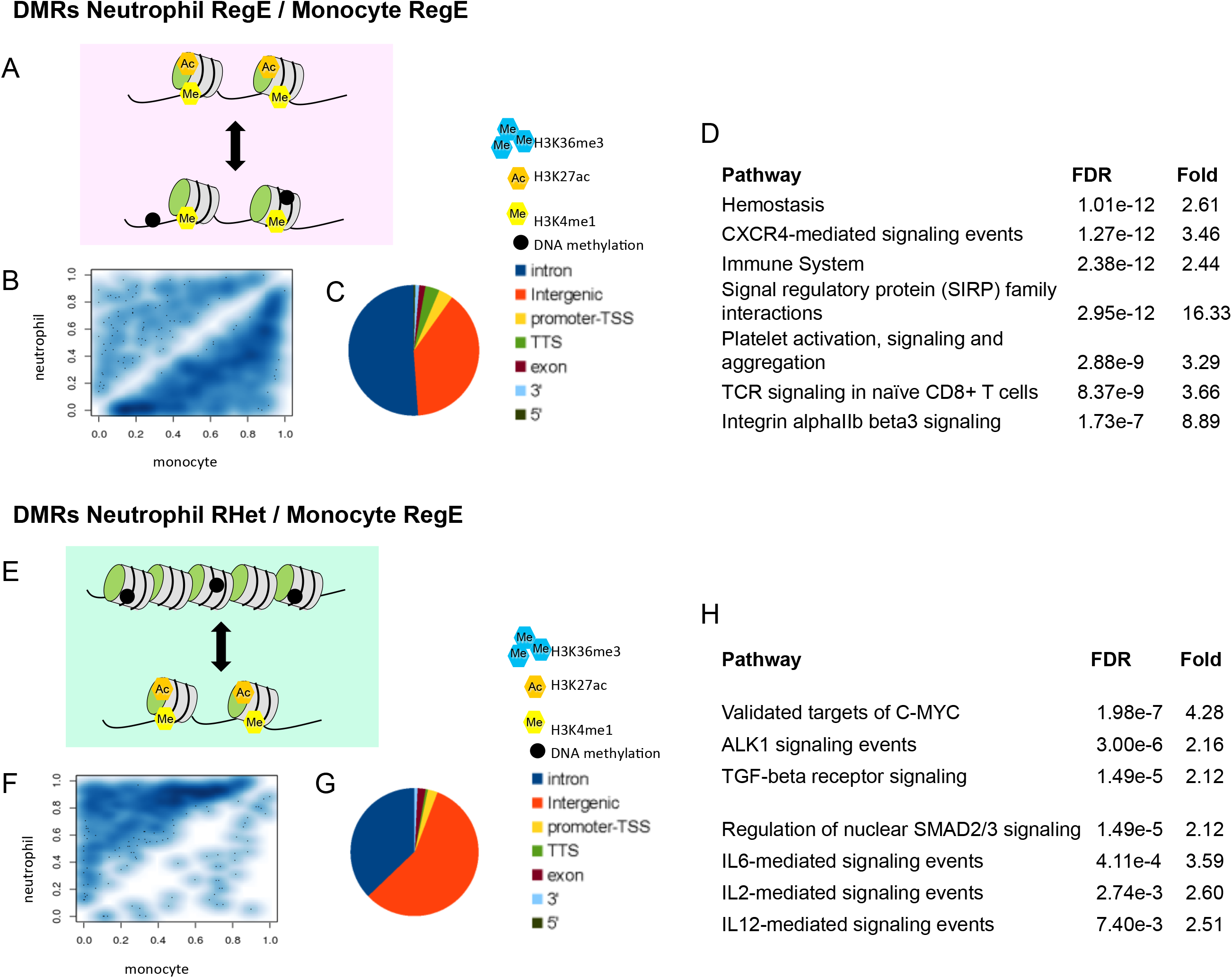
DNA methylation at chromatin segments in neutrophils and monocytes. A-H. Differentially methylated regions (DMRs) are most significantly enriched at common regulatory regions (RegE/RegE, A-D) and at heterochromatic in neutrophils that are monocyte-specific regulatory regions (RHet/RegE, E-H). B, F. Mean methylation in neutrophils and monocytes is compared for each DMR. C, G. Genomic annotation of the DMRs. D, H. Pathway enriched in each DMR group.

These results suggests that the cell-type specific differences observed in methylation of transcribed and regulatory regions were larger than those observed at the promoter regions. The hypermethylated heterochromatic regions in neutrophils that are marked as hypomethylated regulatory elements in monocytes are associated with genes involved in various signaling pathways (GREAT analysis, FDR < 0.01). Interestingly, gene-sets corresponding to targets of c-Myc were enriched in monocyte-specific RegE regions that show differential methylation (Fig. 3H). Taken together, the observed differences in chromatin states between monocytes and neutrophils would suggest a higher transcriptional activity in monocyte, mediated both in cis- and in trans-by more regulatory elements and more active promoters.

### Connecting the epigenome with gene expression regulation

Our results so far suggest that monocytes may have higher transcriptional activity than neutrophils, given that they show higher number of active enhancers, fewer heterochromatic regions and more promoters that have cell-type specific active marks. We performed RNA-Seq on ribo-depleted RNA and mapped, on average, 150 million, 100 nucleotide paired-end reads per sample (Table S9) and used them to quantify different transcriptional elements (Djebali et al., 2012) as annotated in Gencode (Harrow et al., 2012) (version v15; Table S3). We observed >35% of reads mapped to intronic regions both for neutrophils and monocytes (Table S9), an expected result from the used RNA-seq protocol that preserves immature RNAs. PCA based on gene expression separated the samples according to the different biological classes (Fig 1D).

We observed that monocytes exhibit higher overall transcriptional activity than neutrophils. They express more genes (Table S10 and Fig. 4B) and at higher levels (Fig. 4A). We detected 4,334 genes that were differentially expressed between neutrophils and monocytes in adult peripheral blood (FDR < 0.01, and a fold change |logFC| > 2). Consistent with a higher transcriptional activity in monocytes, more than twice as many protein-coding genes and pseudogenes were up-regulated in monocytes compared to neutrophils (Table S11). To try to control for the effect of sequencing coverage when performing RNAseq, we compared the number of detected genes in each sample by serial *‘in silico* dilutions’. We randomly selected subsets of reads and counted the number of genes using an arbitrary threshold of RPKM (reads per kilobase per million mapped reads) > 0.1. We found a read depth independent difference between the number of genes expressed in monocytes and neutrophils, with the latter consistently showing a lower number in cord blood and adult peripheral blood (Fig. 4B). To corroborate if the higher number of genes in monocytes is reflected in higher levels of RNA, we compared the yields of RNA extraction for monocytes and neutrophils from peripheral blood of 48 individuals (reference). We observed that the average RNA quantity extracted from monocytes is 10 times higher than in neutrophils (p < 10^−16^, T-test, Fig. S16).

**Fig. 4.**
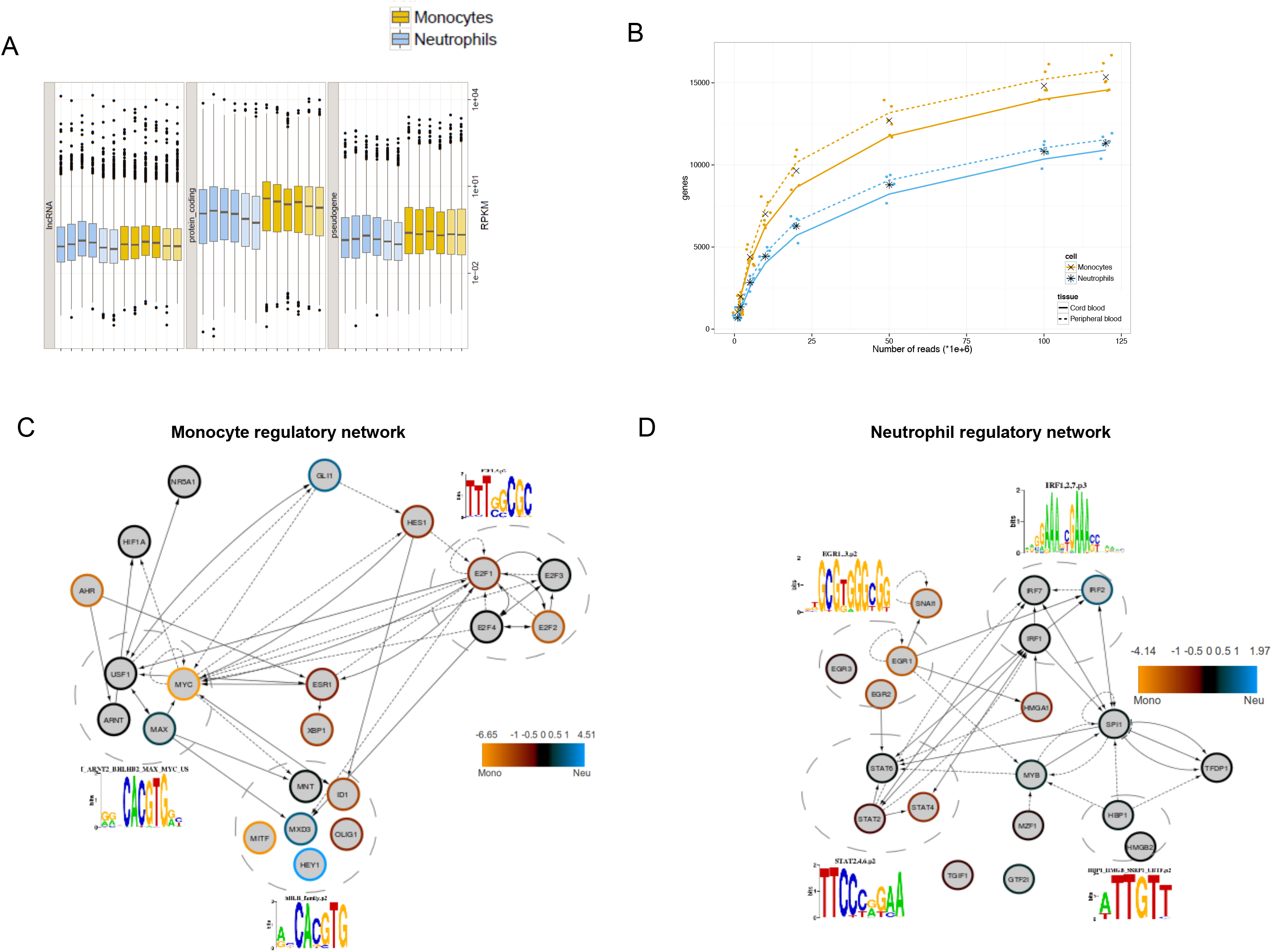
Differential expression in monocytes and neutrophils. A. Boxplots showing expression (RPKM) of lncRNAs, protein coding and pseudogenes in neutrophils (blue) and monocytes (yellow) from peripheral (dark) or cord (light) blood. B. Number of identified genes with an RPKM >0.1 when increasing tag number by random sampling of reads. C-D. Subnetworks of transcription factors identified as putative regulators by ISMARA to be most active in monocytes (C) and neutrophils (D). The networks represent regulatory interactions between the TFs as annotated in TFactS that show expression levels that pass the edgeR filtering for differential expression (see Methods). Dashed circles group TFs that can potentially bind the same motif, represented next to them. Border color of the nodes represents log fold change of expression between monocyte and neutrophil. TFs up-regulated in monocytes (bright yellow) and TFs up-regulated in neutrophils (light-blue).

### Inferring master regulators in monocytes and neutrophils

Having observed a substantial consistency between functional characterization of monocytes and neutrophils at the epigenomic and gene expression level, we proceeded to infer the main regulators in the two cell types using ISMARA (Integrated System for Motif Activity Response Analysis; Piotr at el., 2014). ISMARA identifies the key transcription factors and miRNAs that could drive the observed expression changes between different conditions, in this case our two cell types. ISMARA analysis is focused on known transcription start sites defined with CAGE (Cap Analysis of Gene Expression, Piotr at el., 2014). When we applied it to our RNA-seq data, it identified a total of 31 motifs with a Z-score higher than 2). Of these, 12 are bound by TFs that are predicted to be more active in monocytes and 19 motifs in neutrophils. We integrated the putative regulators identified by ISMARA into networks with their regulatory relationships as annotated in TFactS (Essaghir et al., 2010) and within the context of known functional interactions from REACTOME (Croft et al., 2013). This leads to monocyte- and neutrophil-specific interaction networks (Fig. 4C-D). As shown, many of the master regulators identified in one specific cell-type show higher expression in that cell type.

The most significant motifs predicted by ISMARA as enriched in monocytes are predicted to be bound by the family of basic-helix-loop-helix (bHLH) proteins. Amongst these, based on differential expression, AHR and ARNT are prime candidate regulators given their suggested roles in regulating immunological responses and hematopoietic differentiation (Boitano et al., 2010, Gasiewicz et al., 2010). Another bHLH protein expressed to higher levels in monocytes as compared to neutrophils is MYC that we have shown that is enriched in monocyte-specific enhancers. Finally, E2F motifs are enriched in regulatory regions of monocyte genes, possibly reflecting that the fact that only monocytes retain some proliferative capacity (Dale et al., 2008).

The transcriptional subnetwork that is predicted to be more active in monocytes (1173 genes) is enriched for functions related to multi-cellular organismal processes, regulation of transcription and RNA metabolsim, defense and inflammatory responses. Most of these functions are mediated by MYC targets (553 genes), which dominate the subnetwork. The neutrophil specific transcriptional subnetwork is much smaller than the monocyte one (248 genes) and is not enriched for any specific function.

In neutrophils, ISMARA finds specific enrichment of motifs of the immediate early response gene family (EGR1, 2 and 3) thought to underlie the ability of neutrophils to respond rapidly to inflammatory stimuli (Cullen et al., 2010). In addition, we detect enrichment for motifs bound by factors such as HBP1, HMGA1 and 2, which are architectural elements of chromatin and are involved in the regulation of multiple DNA-dependent processes. In this case, these factors might be potential contributors to the unique spatial organization of neutrophil DNA in a segmented nucleus. Finally, we identified enrichment for motifs bound by factors involved in Interferon and Interleukin response such as the STAT (STAT2,4,6) and the IRF (IRF1, 2 and 7) family. RNA-seq analysis (Fig. S17) revealed a larger number of genes of the type I interferon (IFN) signaling pathway expressed in neutrophils suggesting this to be a specific activated function in neutrophils compared to monocytes. Comparing adult with cord blood neutrophils revealed increased expression of IFN genes in adult blood, potentially reflecting a poised IFN state. Interestingly, despite higher expression of IFN pathway genes in neutrophils, most TSSs in both cell types are marked with active chromatin segments (Fig. S17). These results suggest that, compared to the naive state of cord blood neutrophils, adult circulating neutrophils are already in a more active state, potentially due to previous environmentally induced triggers, the presence of a normal gut flora (which is absent in newborns at the moment of delivery), colonization of the skin and/or ageing.

### Enhancer activity could account for higher gene expression levels in monocytes

We proceeded to investigate which of the epigenomic features mapped contributes most in determining the gene expression in a specific cell-type. Considering factors that are likely to affect gene expression, we focused on DNA methylation and chromatin state at the TSS and on the presence of neighboring active enhancers associated to a specific gene. We divided genes into 100 classes of increasing expression to examine the dependence between the three epigenetic variables and expression. We observe a clear negative correlation in both monocytes and neutrophils between TSS methylation and gene expression but only in the low range of expression (Fig. 5A). This is consistent with our previous observation that DNA methylation at the TSS does not seem to differ substantially between monocytes and neutrophils (Fig. S15).

**Fig. 5.**
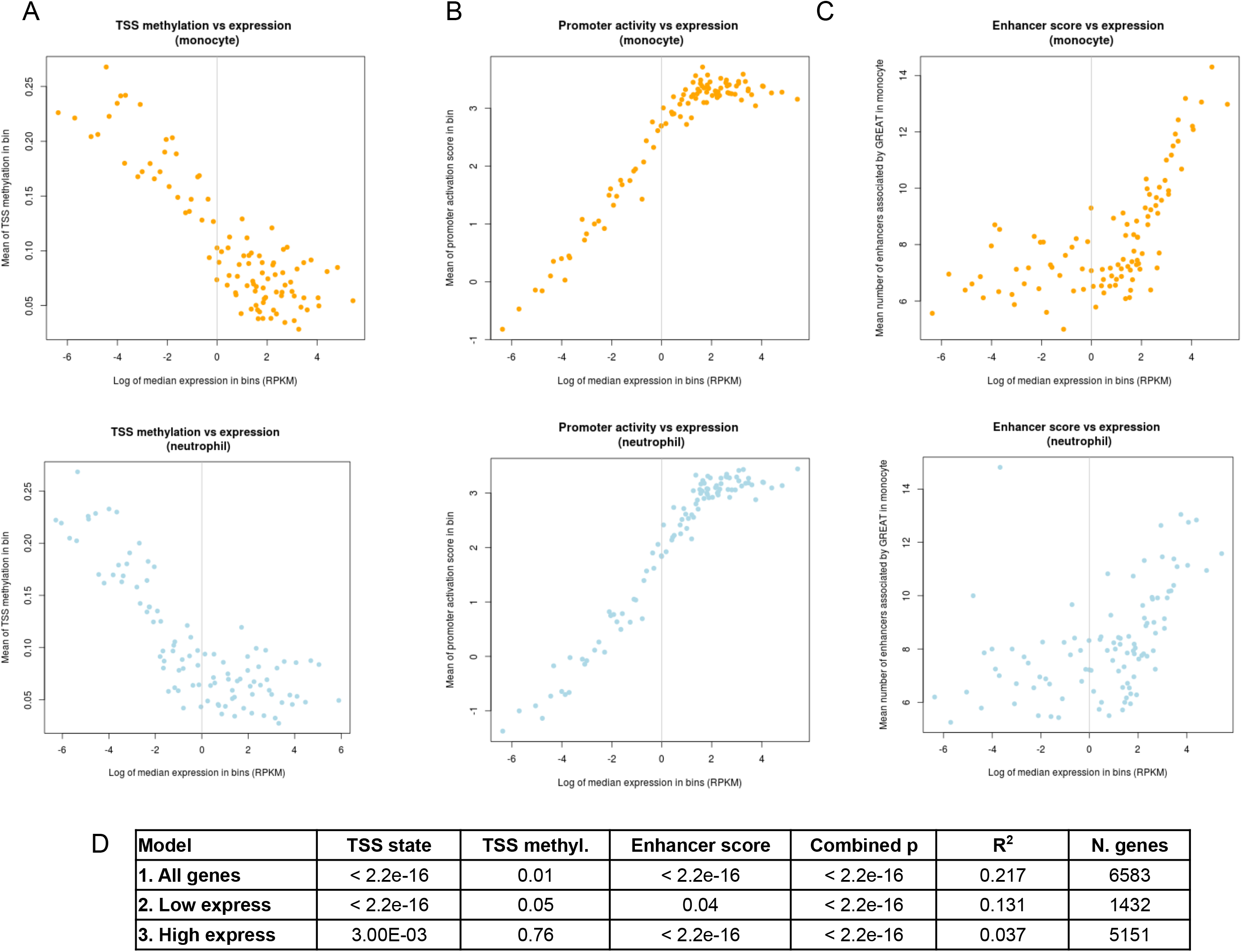
Gene expression dependence on TSS activity (based on histone modification presence), TSS DNA methylation and enhancer score. A. Dependence of monocyte (top) or neutrophil (bottom) gene expression on TSS methylation, represented for 100 classes of increasing gene expression. B. Dependence of monocyte (top) or neutrophil (bottom) gene expression on promoter activity state, represented for 100 classes of increasing gene expression. C. Dependence of monocyte (top) or neutrophil (bottom) gene expression on enhancer score, represented for 100 classes of increasing gene expression. D. Linear models were fit to quantify the relative contribution of each of the three types of epigenetic features (TSS state, TSS DNA methylation and enhancer score) to gene expression in monocytes. Different contribution of each epigenetic feature was observed when fitting separate models for all genes, low-expressed genes (RPKM < 1) and highly-expressed genes (RPKM > 1).

On the contrary, we find a strong positive correlation, persisting into higher expression ranges, between the expression and the promoter activity state (related to the number of samples with active or repressed chromatin state at the promoter in a specific cell-type, see Supp. methods) (Fig. 5B). This result suggest that the histone marks summarised by the chromatin states may have a stronger connection to gene expression than the TSS DNA methylation levels. Finally, we investigated to what extent the cell-type specific gene expression programmes could be influenced by the presence of non-promoter regulatory elements. We defined an *enhancer score* as the number of RegE regions that were associated to each gene by GREAT (McLean et al., 2010), either specifically in one cell type or in both. We observe a strong positive correlation between expression values and the enhancer score, especially in more highly expressed genes (Fig. 5C). Moreover, for the almost 2,000 genes that have enhancer regions associated in both cell types, monocytes show clearly higher enhancer scores (data not shown), suggesting that the presence of more distal regulatory elements in monocytes is an important contribution to the higher transcriptional activity observed in this cell-type.

Our results have shown that the correlation of these three epigenomics features with gene expression varies depending on the range of gene expression levels that we investigate. To assay the statistical significance of the contribution of each epigenomic feature in determining gene expression, we fitted linear regression models using these variables in different combinations (Table S12). As similar results were obtained using the data from neutrophils or from monocytes, we will focus here in discussing the models for monocytes. The most informative factor appears to be the promoter activity state, explaining by itself 19% of the variance in expression. (Table S12). Adding the enhancer score to the model further improves the results (21.6% of variance explained) while it remains unchanged by the addition of the TSS methylation in the full model (21.7% of variance explained).

We then built three different multivariate models with the three epigenomic features: one model for all genes, a second model for low expressed genes, and a third model for high expressed genes (Fig. 5D). TSS DNA methylation only has a significant contribution for those genes that are expressed at low levels (p = 0.05) but it is not a relevant feature for highly expressed genes (p = 0.76). Promoter marking as the main predictor of gene activity of low expressed genes (p < 10^−16^), while enhancer activity is most predictive for higher expressed genes (p < 10^−16^). These results indicate that the number of enhancers per gene can be an important factor regulating the expression levels of many genes.

## DISCUSSION

Knowledge of chromatin organization is fundamental to our understanding of how different cell types arise from a single hematopoietic stem cell. Although in recent years many epigenomic datasets have become available, most notably through the Encyclopedia of DNA Elements (ENCODE) project (http://www.genome.gov/10005107), these datasets have been largely restricted to immortalized cell lines. In contrast, comprehensive epigenomic analysis of primary human cell types has only been initiated in recent years (Satterlee et al., 2010; Bernstein et al., 2010). In this manuscript, BLUEPRINT generated and analyzed the first reference epigenomic data sets of primary human neutrophils and monocytes. For a total of 12 samples we describe the detailed and integrated analysis of RNA-seq data, ChIP-seq of six histone modifications and DNA methylation by bisulfite sequencing at base pair resolution for neutrophils and monocytes isolated from peripheral or cord blood, thereby providing a first insight into the epigenetic programming of two different types of myeloid cells as well as providing a reference for ongoing efforts to examine epigenetic variation within monocytes and neutrophils obtained from 200 healthy individuals.

Genome segmentation based on histone modifications allowed the identification of genomic elements with different chromatin states in the two cell types. The chromatin states identified with the combination of these histone marks are similar to previously reported chromatin segments in cell lines and other primary cells (Hoffman et al., 2013). Remarkably, in comparison to neutrophils, monocytes have more regulatory regions that are active, probably reflecting a more plastic epigenome that can still undergo further differentiation into several types of macrophages (Saeed et al., submitted), dendritic cells and osteoclasts. In contrast, most of the neutrophil-specific features tend to be in a silent chromatin state. However, it should be noted that the majority of the heterochromatin is characterized by the absence of signal (state 2). Therefore, we cannot rule out the possibility that part of these neutrophil-specific ‘heterochromatin’ segments may be decondensed chromatin (Martinod et al., 2013).

Our analysis of differential DNA methylation in the context of chromatin states allowed us to identify regulatory regions (enhancers) as the most frequent chromatin regions were differential methylation occurs. Most monocyte-specific regulatory regions were hypomethylated compared with the heterochromatic-like and high methylation status of these regions in neutrophils. TSS methylation does not seem to differ substantially between the two cell types and is unlikely to be mediating the regulation of the main gene expression programs that drive the different functions of monocytes and neutrophils. Indeed, modeling of epigenetic features related to gene expression correlates mostly with the presence of putative enhancers and marks of active promoters in the TSS of the corresponding genes.

Although a subset of genes is more highly expressed in neutrophils (such as many of the IFN pathway genes), our analysis revealed generally higher transcriptional activity in monocytes. More genes with high RPKM values were detected in monocytes, while in neutrophils relatively more genes had lower RPKM values. Accordingly, monocytes yield higher levels of RNA, further strengthening the notion that in monocytes genes are generally higher expressed. In monocytes more genes are associated with multiple regulatory elements, which likely represent super-enhancers (Hnisz et al., 2013) that reflect the cells identity. Interestingly, one of these enhancer clusters is associated with the transcription factor MYC which is higher expressed in monocytes than neutrophils and has been described as a transcriptional amplifier (Nie, 2012). We found that MYC binding motif is enriched at monocyte-specific enhancers and that DMRs are enriched in known targets of MYC. In an independent analysis where epigenomic information was not used, ISMARA predicted that the gene expression of MYC targets could be associated to a higher activity of MYC in monocytes. It is thus tempting to speculate that the higher MYC levels in monocytes might have an important role for the higher transcription in monocytes compared to neutrophils. Finally, we examined the relative contribution of each epigenetic mark to gene expression. Interestingly, the number of enhancers associated per gene is the most informative feature to predict gene expression levels of highly expressed genes. These simple models suggest that the highest number of enhancers in monocytes could indeed play a role in their higher gene expression levels.

Altogether, our study provides comprehensive epigenetic charts of chromatin states in primary human phagocytes as well as new insights into the regulatory program of neutrophils and monocytes, thus providing a comprehensive resource that can be used for studying the regulation of the human innate immune system.

## ACKNOWLEDGEMENTS

The work described in the manuscript was supported by the European Union’s Seventh Framework Program through the BLUEPRINT grant with code HEALTH-F5-2011-282510. NS’s research was further supported by the Wellcome Trust (Grant Codes WT098051 and WT091310), the EU FP7 (EPIGENESYS Grant Code 257082 and BLUEPRINT Grant Code HEALTH-F5-2011-282510), research in the Ouwehand laboratory is further supported by program grants from the National Institute for Health Research (to WHO, NF; http://www.nihr.ac.uk) and the British Heart Foundation (to AR; http://www.bhf.org.uk) under numbers RP-PG-0310-1002 and RG/09/12/28096. AV was also funded by Spanish Ministry of Economy and Competitivity (MINECO, BIO2012-40205). Both the Cambridge BioResource and FACS cell sorting facility were supported by a NIHR grant to the Cambridge NIHR Biomedical Research Centre. VP was supported by a FEBS long-term fellowship. DR’s research is supported by a Wellcome Trust Seed Award in Science (206103/Z/17/Z).

## SUPPLEMENTARY METHODS

### Cord blood collection

Cord blood was collected with informed consent (REC 12/EE/0040) at the Rosie Hospital, Cambridge University Hospitals, Cambridge, UK and processed within 18 hours.

### Peripheral adult blood collection

Peripheral blood was collected with informed consent (ReC 12/EE/0040) at NHS Blood and Transplant in Cambridge, from volunteers belonging to the NHSBT Cambridge BioResource (http://www.cambridgebioresource.org.uk/) and processed within 2 hours.

### Isolation of neutrophils and monocytes

A whole unit (460 ml) of peripheral blood or a large unit (>120 grams) of cord blood were used. Diluted blood was separated by gradient centrifugation (Percoll 1.078 g/ml) and neutrophils were isolated from the pellet, after red blood cell lysis, by CD16 positive selection (Miltenyi). Leukocytes were further fractionated to obtain a monocyte rich layer using a second gradient (Percoll 1.066 g/ml). Monocytes were purified using CD16 depletion and subsequent a positive selection for the CD14-positive monocytes (Miltenyi). The purity of each preparation was assessed by flow cytometry on a Beckman Coulter FC500, by expression with Illumina HT-12v4 arrays (E-MTAB-1573 at arrayexpress) and finally by microscopic inspection of morphology using appropriate stained cytospin preparations. Samples which did not meet predefined criteria of cell purity were not included in the analysis. The purified cells were processed to generate genomic DNA for WGBS, RNA (Trizol extraction according to manufacturer’s instructions; Agilent BioAnalyzer RIN >8) and cells were fixed for ChIP in 1% formaldehyde. Detailed versions of all protocols are available at http://www.blueprint-epigenome.eu/ under the result section. Representative pictures of monocytes and neutrophils are shown in Supplementary figure 1.

### Chip-seq data production

Antibodies for H3K4me1 (H3 lysine 4 monomethylation), H3K4me3 (H3 lysine 4 trimethylation), H3K9me3, H3K27me3, H3K36me3 and H3K27ac (H3 lysine 27 acetylation). were extensively characterized and experimental procedures and analyses were optimized for the two cell types (www.blueprint-epigenome.eu). We mapped, on average, 40 million, 50 nucleotide single-end reads per ChIP-seq sample using BWA (Li et al., 2009).

### Peak Calling

For peak calling the BAM files were first filtered to remove reads with mapping quality less than 15, followed by modelling of fragment size (http://code.google.com/p/phantompeakqualtools/). Peak calling algorithm MACS2 (http://github.com/taoliu/MACS/) was used to detect the binding sites for the six histone marks in study at FDR (q-value) 0.05. Histone marks such as H3K27me3, H3K36me3, H3K9me3, and H3K4me1 were called at *broad* setting of MACS2 while the other two H3K27ac and H3K4me3 were called at default (*narrow*) setting.

### Differential Peaks

To identify differential binding sites, tags in binding sites of histone marks were counted in neutrophils and monocytes, followed by application of DESeq algorithm (Anders & Huber, 2010).

**Coverage Profile**. NGSplot (http://code.google.com/p/ngsplot/) was used to get the distribution of different histone marks over the genebody.

### DNasel-seq

DNaseI libraries were prepared as described (John et al. 2013). In brief nuclei were isolated from monocytes using Buffer A (15 mM NaCl, 60mM KCl, 1 mM EDTA, pH 8.0,0.5 mM EGTA, pH 8.0, 15 mM Tris-HCl, pH 8.0, 0.5 mM Spermidine) supplemented with 0.015% IGEPAL CA-630 detergent. DNaseI treatment was done for 3 minutes and the reaction was stopped with stop buffer (50 mM Tris-HCl, pH 8, 100 mM NaCl, 0.10% SDS, 100 mM EDTA, pH 8.0, 1 mM Spermidine, 0.3 mM Spermine) The sample was further fractionated on 9% Sucrose gradient for 24hrs/25000 rpm at 16°C. Fractions of less than 1kb fragments were purified and used as input material for the illumina library preparation protocol.

Calling accessibility hotspots (DNase-I Hypersensitive Sites, DHSs) identified 130,000-180,000 high confidence DHSs, which are for the most part located outside of promoter regions (Fig S5B, C). PCA analysis as well as hierarchical clustering analysis of our dataset together with ENCODE DNaseI-seq profiles of other blood cell types, revealed a clear cell type driven clustering of the monocyte DNAseI-seq tracks (data not shown), showing the reproducibility of the applied technology. To obtain high sequencing depth and due to the high similarity, we pooled the four monocytes DNase-seq data sets. Further footprinting detection was performed using the pipeline as described (Neph et al., 2012).

### Genome segmentation

We used ChIP-seq data to segment the genome in different chromatin states depending on the combination of different histone modifications. For this a multivariant Hidden Markov Models (HMM) was used. This model uses two types of information, the frequency with which different chromatin mark combinations are found with each other and the frequency with which different chromatin states occur in spatial relationships of each other along the genome. To apply this method we used the implementation as described by Ernst et al. in ChromHmm software (v1.03). The input data to generate the model were the ChIP-seq bed files containing the genomic coordinates and strand orientation of mapped sequences (after removal of duplicate reads). First the genome was divided in 200 bp non-overlapping intervals which we independently asigned if each of the 6 chromatin modifications marks was detected as present (1) or not (0) based on the count of tags mapping to the interval and on the basis of a Poisson background model using a threshold of 10^−4^ as explained in Ernst et al. In those samples where a chromatin modification profile was missing, a missing value for the interval was assigned (2).

After binarization of each chromatin modification mark for each sample we used all the cell type samples to train the HMM model using a fixed number of randomly-initialized hidden states, varying from 9 up to 13 states. We focused on a 11-state model that provides sufficient resolution to resolve biologically meaningful chromatin patterns. We used this model to compute the probability that each location is in a given chromatin state, and then assigned each 200-bp interval to its most likely state for each sample. Consecutive intervals within the same chromatin state were joined and the length statistics for each chromatin state were calculated in R (v2.14.1). In those samples (C000S5, C0010K, C001UY, all cases are monocytes) that a chromatin modification ChIP-seq was missing the following imputation in the 200 bp intervals of the binarized files was carried out; 0 when in the other 4 samples from the same cell type the mark is absent; 1 when at least in 3 out of 4 samples the mark is present; 2 (missing) all the cases not considered as 0 or 1.

The percentage of genome overlap for each state and different annotation data was computed with ChromHmm software (v1.03) as described by Ernst et al. The gene and miRNA annotations were from the Genecode Project (http://www.gencodegenes.org) version 15 (01/2013) based on GRC37/hg19. The sequence data for CpG islands, repeats and nuclear lamina were obtained from the UCSC genome browser. The DnaseI and Hyper/Hypo-methylated regions were obtained as described in Supp Methods.

### Differential chromatin states between cell types

For each cell type, the chromatin state assigment at each 200bp interval was compared. We calculated the frequency that a state appeared in the same interval for each cell type. Intervals where all samples shared the same state in one cell type but shared a different state in the other cell type were defined as intervals with differential chromatin states.

### Functional enrichment analysis

Functional enrichment of enhancers and DMRs were carried out with GREAT software v2.0.2 (McLean, 2010) using the default parameters in the web server (http://bejerano.stanford.edu/great).

### Genomic segments annotation

Genomic annotation of enhancer and DMR segments were carried out with Hypergeometric Optimization of Motif EnRichment (HOMER) software v4.2 (Heinz S et al, 2010). The tool annotatePeaks.pl was used with parameters by default and defined in the help. A gtf file from the Genecode Project (http://www.gencodegenes.org) version 15 (01/2013) based on GRC37/hg19 was used for annotations. The annotation includes wether a segment is in the TSS (transcription start site), TTS (transcription termination site), Exon (Coding), 5’ UTR Exon, 3’ UTR Exon, Intronic, or Intergenic. Since some annotation overlap, a priority is assign based on the following: 1. TSS (by default defined from −1kb to +100bp); 2. TTS (by default defined from −100 bp to +1kb); 3. CDS Exons; 4. 5′ UTR Exons; 5. 3′ UTR Exons; 6. Introns; 7. Intergenic. More detailed information in http://homer.salk.edu/homer/ngs/annotation.html

### Motif enrichment analysis in enhancer regions

Motif analysis for monocyte-specific and neutrophil-specific enhancer regions were carried out with Hypergeometric Optimization of Motif EnRichment (HOMER) software v4.2 (Heinz S et al, 2010). The tool findMotifs.pl was used with enhancer regions in fasta format and parameters by default and defined in the help. The background used was the common enhancer regions plus monocyte-specific or neutrophil-specific enhacer regions for each analysis.

### Motif actitivity response analysis

The motif activity response analysis were carried out with ISMARA (Integrated System for Motif Actitivity Response Analysis) software accessed by webserver May 2014. ISMARA predicts regulatory sites for transcription factors (TFs) and micro-RNAS (miRNAs) from RNA-seq driving gene expression changes across cell types. A Z-score, which summarizes the importance of the motif for explaining the expression variation across cell types, higher than 2 was considered for significant motifs.

### Whole-genome bisulfite sequencing and library construction

Genomic DNA (1-2μg) was spiked with unmethylated λ DNA (5ng of λ DNA per μg of genomic DNA) (Promega). The DNA was sheared by sonication to 50-500bp using a Covaris E220 and fragments of size 150-300 bp were selected using AMPure XP beads (Agencourt Bioscience Corp.). Genomic DNA libraries were constructed using the Illumina TruSeq Sample Preparation kit (Illumina Inc.) following the lllumina standard protocol: end repair was performed on the DNA fragments, an adenine was added to the 3’ extremities of the fragments and Illumina TruSeq adapters were ligated at each extremity. Adter adaptor ligation, the DNA was treated with sodium bisulfite using the EpiTexy Bisulfite kit (Qiagen) following the manufacturer’s instructions for formalin-fixed and paraffin-embedded (FFPE) tissue samples. Two rounds of bisulfite conversion were performed to assure a high conversion rate. An enrichment for adaptor-ligated DNA was carried out through 7 PCR cycles using the PfuTurboCx Hotstart DNA polymerase (Stratagene). Library quality was monitored using the Agilent 2100 BioAnalyzer (Agilent), and the concentration of viable sequencing fragments (molecules carrying adaptors at both extremities) estimated using quantitative PCR with the library quantification kit from KAPA Biosystem. Paired-end DNA sequencing (2x100 nucleotides) was then performed using the Illumina Hi-Seq 2000. Amounts of sequence reads and the proportion of aligned reads are shown in Supplementary Table S7.

### Read mapping and estimation of cytosine methylation levels

Read mapping was carried out using the GEM aligner (v1.242) against a composite reference containing two copies of the human GRCh37 reference and two copies of the NCBI viral genome database (v35). For both the human and viral references, one copy had all C bases replaced by T and the other had all G bases replaced by A. The names of the contigs in the combined reference FASTA file were modified by adding #C2T or #G2A to the end of the contig names depending on the conversion performed. Before mapping was performed the original sequence of each read was stored. The first read of each pair then had all C bases replaced by T, and the second read had all G bases replaced by A. Read mapping with GEM was performed allowing up to 4 mismatches per read from the reference. After read mapping the original sequence of each read was restored.

Estimation of cytosine levels was carried out on read pairs where both members of the read mapped to the same contig with consistent orientation, and there was no other such configuration at the same or less edit distance from the reference. After mapping, we restored the original read data in preparation for the inference of genotype and methylation status. We estimated genotype and DNA methylation status simultaneously using software developed at the Centro Nacional de Análisis Genómico, taking into account the observed bases, base quality scores and the strand origin of each read pair. For each genomic position, we produced estimates of the most likely genotype and the methylation proportion (for genotypes containing a C on either strand). A phred scaled likelihood ratio for the confidence in the genotype call was estimated for the called genotype at each position. For each sample, CpG sites were selected where both bases were called as homozygous CC followed by GG with a Phred score of at least 20, corresponding to an estimated genotype error level of <=1%. Sites with >500x coverage depth were excluded to avoid centromeric/telomeric repetive regions. A common set of called CpG sites for all analyzed samples was generated, and all subsequent analyses used this common set.

### Differentially methylated regions (DMRs)

To detect differential methylation at chromatin segments RnBeads (Assenov et al., 2013), an R package for comprehensive analysis of DNA methylation data obtained with bisulfite sequencing protocols, was used.

### Genomic annotation of CpG sites

CpG sites in the selected common set were annotated using data from the version 15 of the Gencode annotation database. For the location relative to a gene, the following categories were used: TSS 1500 (from 201 to 1,500 bp upstream of the transcriptional start site (TSS)), TSS 200 (from 1 to 200 bp upstream of the TSS), 5 UTR, first exon, gene body (from the first intron to the last exon), 3 UTR and intergenic regions. For the location relative to a CpG island (CGI), the following groups were used: within CGI, in CGI shore (0–2 kb from the CGI edge), in CGI shelf (>2 kb to 4 kb from the CGI edge) and outside CGI. Owing to the presence of alternative transcription start sites and regions containing more than one gene, some of the CpGs were assigned multiple annotations.

### RNA-seq data processing

The raw reads were aligned to the reference genome GRCh37 with GEMtools (version 1.6.2, ref::gemtools). The mapped reads were used to quantify exons, splice junctions, transcripts and genes with the Flux Capacitor (version 1.2.4; Montgomery et al., 2010), as annotated in GENCODE version 15 (Harrow et al., 2012). The expression levels were quantified using RPKM, as previously described (Mortazavi et al., 2008). Using GEMtools, we mapped on average 150 million, 100 nucleotide paired-end reads per sample.

### Differential gene expression analysis

The differential gene expression analysis was performed with the R package edgeR (version 3.0.8; Robinson et al., 2010). Following the package instruction, we used read counts for all the genes tested. We searched for differentially expressed genes in all the pairwise combinations of tissues and cell types. The initial set included 47,507 genes, excluding the small RNAs. A different number of genes in all the comparisons was removed before analysis to filter for lowly expressed genes (Robinson et al., 2010). Only those genes with a cpm (count per million reads) >= 1 in at least two samples were finally tested.

### Exonic vs total ratio

For each gene, excluding intronless genes, we counted the number of reads mapping to exonic and purely intronic regions. A purely intronic region is defined as a genic region that does not overlap any exon on the same strand. Exonic and intronic read counts were normalized by the total number of exonic and purely intronic nucleotides, respectively, in each gene. Finally, the ratio between the normalized exonic read counts, and the total normalized read counts (exonic plus intronic). To filter for possible noise, we considered only the genes with a total number of read counts > 0.01.

### Statistical models of expression regulation

Models of gene expression were generated using the following measurements for monocyte samples:

Promoter activity state. A score ranging from -4 to 4 was assigned to each promoter based on the chromatin state assigned to the segment. For each gene and each sample the chromatin state ‘Active promoter’ (Apro) or ‘Repressed promoter’ (Rpro) was annotated. The number of samples with Apro and Rpro marks were counted and their difference was calculated. Since there are 4 samples for each cell type the maximum activation of a promoter is found when all 4 samples have Apro state (promoter acrivity score=+4), whereas the maximum repression of a promoter is found when all 4 samples have Rpro state (promoter activity score=−4), with numbers in between indicating discordance between the 4 samples.

TSS methylation. Methylation Beta values were taken for the TSS 200 defined segment for each gene and the median was taken across the 4 samples.

Enhancer score. The enhancer score was defined based on the number of regulatory regions assigned to each gene by GREAT. For example, to calculate the score for genes that presented association with regulatory regions in multiple cell-types the number of regulatory regions associated to a gene specifically in monocite were summed with the ones that are present in both cell types. For the genes that have regulatory regions associated to them only in monocyte only that number was considered. The score was calculated for all genes that are associated to regulatory regions in any cell type, such that they can have a score of 0 in the case in which in a specific cell type there are no associations between that gene and regulatory regions. This score is thus a discrete numerical value ranging from 0 to the maximum number of regions that associate to a specific promoter (84 in monocytes and 67 in neutrophils).

Gene expression binning. Genes were subdivided into 100 classes of gene expression (each containing 139 to 140 genes), based on the RPKM median values across the 4 samples in a specific cell-type. For each class of expression the median values of the above defined variables were calculated and plotted against the median values of gene expression.

Linear regression models. Having observed clear linear dependencies between the promoter activity state score, enhancer score, methylation at the TSS and gene expression, we constructed a linear regression model with these variables (See Table S12, function lm in R). Since we observed a change in the trend of expression as a function of each variable between genes with RPKM above and below 1, we decided to train separate models for these two classes of genes, running the regression on subsets of the data.

### Blueprint Data Availability

The Blueprint samples were consented for managed access release that means that the analysis results we present in this paper fall under two categories: Unique data, which represents the sequence, alignment and genotype calls, and non Unique data that represents the signal, quantification and methylation states across the genome for our different samples.

Unique data is only available by consent of the Blueprint Data Access Committee through the European Genome Phenome Archive (EGA) (http://www.ega.ac.uk). Details on how to apply for access can be found on our website. (http://www.blueprint-epigenome.eu/index.cfm?p=B5E93EE0-09E2-5736-A708817C27EF2DB7) The EGA Data set identifiers are listed in Table S8. The non-unique data is available through numerous routes as described in Table S9. Our ftp site provides both flat files in appropriate formats (Table S10) and a trackhub that allows our files to be attached to both UCSC (Karolchik et al., 2014) and Ensembl (Flicek et al., 2014) browsers in one go rather than one at a time. We also present data from the Blueprint project in a BioMart interface to allow easy data mining and a Genomatix browser. A complete list of the raw files available from the ftp is listed together with associated meta data in the data index file. (ftp://ftp.ebi.ac.uk/pub/databases/blueprint/releases/20130617/homo_sapiens/20130617.data.index)This file lists all the files associated with the non unique primary analysis and details like cell type, sample supplier and disease status. A full list of columns is given in Table S11. This paper also collects together several higher level analyses on these data types. These can all be found in the directory ftp://ftp.ebi.ac.uk/pub/databases/blueprint/paper_data_sets/monocyte_neutrophil_2014http://ftp.ebi.ac.uk/pub/databases/blueprint/paper_data_sets/monocyte_neutrophil_2014 The contents of the directory itself is described in Table S3.

### Supplementary websites

Gemtools: https://github.com/gemtools/gemtoolshttps://github.com/gemtools/gemtools https://github.com/gemtools/gemtools

Webflux: http://sammeth.net/confluence/display/FLUX/Homehttp://sammeth.net/confluence/display/FLUX/Home

## REFERENCES

Abbott, A. (2011). Europe to map the human epigenome. Nature 477, 518.

Adams, D., Altucci, L., Antonarakis, S.E., Ballesteros, J., Beck, S., Bird, A., Bock, C., Boehm, B., Campo, E., Caricasole, A., et al. (2012). BLUEPRINT to decode the epigenetic signature written in blood. Nat Biotechnol 30, 224–226.

Adcock, I.M. (2007). HDAC inhibitors as anti-inflammatory agents. Br J Pharmacol 150, 829–831.

Anders, S., and Huber, W. (2010). Differential expression analysis for sequence count data. Genome Biol 11, R106.

Andersson, R., Gebhard, C., Miguel-Escalada, I., Hoof, I., Bornholdt, J., Boyd, M., Chen, Y., Zhao, X., Schmidl, C., Suzuki, T., et al. (2014). An atlas of active enhancers across human cell types and tissues. Nature 507, 455–461.

Assenov Y, Müller F, Lutsik P, Walter J, Lengauer T, Bock C. Nat Methods. 2014Nov;11(11):1138–1140.

Balwierz, P.J., Pachkov, M., Arnold, P., Gruber, A.J., Zavolan, M., and van Nimwegen, E. (2014). ISMARA: automated modeling of genomic signals as a democracy of regulatory motifs. Genome Res 24, 869–884.

Bernstein, B.E., Stamatoyannopoulos, J.A., Costello, J.F., Ren, B., Milosavljevic, A., Meissner, A., Kellis, M., Marra, M.A., Beaudet, A.L., Ecker, J.R., et al. (2010). The NIH Roadmap Epigenomics Mapping Consortium. Nat Biotechnol 28, 1045–1048.

Blake, J.A., Bult, C.J., Eppig, J.T., Kadin, J.A., and Richardson, J.E. (2009). The Mouse Genome Database genotypes::phenotypes. Nucleic Acids Res 37, D712–719.

Bock, C., Beerman, I., Lien, W.H., Smith, Z.D., Gu, H., Boyle, P., Gnirke, A., Fuchs, E., Rossi, D.J., and Meissner, A. (2012). DNA methylation dynamics during in vivo differentiation of blood and skin stem cells. Mol Cell 47, 633–647.

Boitano AE, Wang J, Romeo R, Bouchez LC, Parker AE, Sutton SE, Walker JR, Flaveny CA, Perdew GH, Denison MS, Schultz PG, Cooke MP (2010). Aryl hydrocarbon receptor antagonists promote the expansion of human hematopoietic stem cells. Science 329 (5997): 1345–8.

Borregaard, N., and Herlin, T. (1982). Energy metabolism of human neutrophils during phagocytosis. J Clin Invest 70, 550–557.

Brinkman AB, Gu H, Bartels SJ, Zhang Y, Matarese F, Simmer F, Marks H, Bock C, Gnirke A, Meissner A, Stunnenberg HG. (2012). Sequential ChIP-bisulfite sequencing enables direct genome-scale investigation of chromatin and DNA methylation cross-talk. Genome Research 22(6):1128–38.

Chacko, B.K., Kramer, P.A., Ravi, S., Johnson, M.S., Hardy, R.W., Ballinger, S.W., and Darley-Usmar, V.M. (2013). Methods for defining distinct bioenergetic profiles in platelets, lymphocytes, monocytes, and neutrophils, and the oxidative burst from human blood. Lab Invest 93, 690–700.

Croft, D., Mundo, A.F., Haw, R., Milacic, M., Weiser, J., Wu, G., Caudy, M., Garapati, P., Gillespie, M., Kamdar, M.R., et al. (2013). The Reactome pathway knowledgebase. Nucleic Acids Res 42, D472–477.

Dale, D.C., Boxer, L., and Liles, W.C. (2008). The phagocytes: neutrophils and monocytes. Blood 112, 935–945.

Devalon, M.L., Elliott, G.R., and Regelmann, W.E. (1987). Oxidative response of human neutrophils, monocytes, and alveolar macrophages induced by unopsonized surface-adherent Staphylococcus aureus. Infect Immun 55, 2398–2403.

Djebali, S., Davis, C.A., Merkel, A., Dobin, A., Lassmann, T., Mortazavi, A., Tanzer, A., Lagarde, J., Lin, W., Schlesinger, F., et al. (2012). Landscape of transcription in human cells. Nature 489, 101–108.

Ernst, J., and Kellis, M. (2010). Discovery and characterization of chromatin states for systematic annotation of the human genome. Nat Biotechnol 28, 817–825.

Essaghir, A., Toffalini, F., Knoops, L., Kallin, A., van Helden, J., and Demoulin, J.B. (2010). Transcription factor regulation can be accurately predicted from the presence of target gene signatures in microarray gene expression data. Nucleic Acids Res 38, e120.

Fairfax, B.P., Humburg, P., Makino, S., Naranbhai, V., Wong, D., Lau, E., Jostins, L., Plant, K., Andrews, R., McGee, C., and Knight, J.C. (2014). Innate immune activity conditions the effect of regulatory variants upon monocyte gene expression. Science 343, 1246949.

Galli, S.J., Borregaard, N., and Wynn, T.A. (2011). Phenotypic and functional plasticity of cells of innate immunity: macrophages, mast cells and neutrophils. Nature immunology 12, 1035–1044.

Gasiewicz TA, Singh KP, Casado FL (March 2010). The aryl hydrocarbon receptor has an important role in the regulation of hematopoiesis: implications for benzene-induced hematopoietic toxicity. Chem. Biol. Interact. 184 (1-2): 246–51

Guelen, L., Pagie, L., Brasset, E., Meuleman, W., Faza, M.B., Talhout, W., Eussen, B.H., de Klein, A., Wessels, L., de Laat, W., and van Steensel, B. (2008). Domain organization of human chromosomes revealed by mapping of nuclear lamina interactions. Nature 453, 948–951.

Harrow, J., Frankish, A., Gonzalez, J.M., Tapanari, E., Diekhans, M., Kokocinski, F., Aken, B.L., Barrell, D., Zadissa, A., Searle, S., et al. (2012). GENCODE: the reference human genome annotation for The ENCODE Project. Genome Res 22, 1760–1774.

Hindorff, L.A., Sethupathy, P., Junkins, H.A., Ramos, E.M., Mehta, J.P., Collins, F.S., and Manolio, T.A. (2009). Potential etiologic and functional implications of genome-wide association loci for human diseases and traits. Proc Natl Acad Sci U S A 106, 9362–9367

Hnisz, D., Abraham, B.J., Lee, T.I., Lau, A., Saint-Andre, V., Sigova, A.A., Hoke, H.A., and Young, R.A. (2013). Super-enhancers in the control of cell identity and disease. Cell 155, 934–947.

Hodges, E., Molaro, A., Dos Santos, C.O., Thekkat, P., Song, Q., Uren, P.J., Park, J., Butler, J., Rafii, S., McCombie, W.R., et al. (2011). Directional DNA methylation changes and complex intermediate states accompany lineage specificity in the adult hematopoietic compartment. Mol Cell 44, 17–28.

Hoffman, M.M., Ernst, J., Wilder, S.P., Kundaje, A., Harris, R.S., Libbrecht, M., Giardine, B., Ellenbogen, P.M., Bilmes, J.A., Birney, E., et al. (2013). Integrative annotation of chromatin elements from ENCODE data. Nucleic Acids Res 41, 827–841.

Ji, H., Ehrlich, L.I., Seita, J., Murakami, P., Doi, A., Lindau, P., Lee, H., Aryee, M.J., Irizarry, R.A., Kim, K., et al. (2010). Comprehensive methylome map of lineage commitment from haematopoietic progenitors. Nature 467, 338–342.

Joshi-Tope, G., Gillespie, M., Vastrik, I., D’Eustachio, P., Schmidt, E., de Bono, B., Jassal, B., Gopinath, G.R., Wu, G.R., Matthews, L., et al. (2005). Reactome: a knowledgebase of biological pathways. Nucleic Acids Res 33, D428–432.

Jostins, L., Ripke, S., Weersma, R.K., Duerr, R.H., McGovern, D.P., Hui, K.Y., Lee, J.C., Schumm, L.P., Sharma, Y., Anderson, C.A., et al. (2012). Host-microbe interactions have shaped the genetic architecture of inflammatory bowel disease. Nature 491, 119–124.

Kieffer-Kwon, K.R., Tang, Z., Mathe, E., Qian, J., Sung, M.H., Li, G., Resch, W., Baek, S., Pruett, N., Grontved, L., et al. (2013). Interactome maps of mouse gene regulatory domains reveal basic principles of transcriptional regulation. Cell 155, 1507–1520.

Kimura, T., Kadokawa, Y., Harada, H., Matsumoto, M., Sato, M., Kashiwazaki, Y., Tarutani, M., Tan, R.S., Takasugi, T., Matsuyama, T., et al. (1996). Essential and non-redundant roles of p48 (ISGF3 gamma) and IRF-1 in both type I and type II interferon responses, as revealed by gene targeting studies. Genes Cells 1, 115–124.

Kolaczkowska, E., and Kubes, P. (2013). Neutrophil recruitment and function in health and inflammation. Nat Rev Immunol 13, 159–175.

Kulis, M., Heath, S., Bibikova, M., Queiros, A.C., Navarro, A., Clot, G., Martinez-Trillos, A., Castellano, G., Brun-Heath, I., Pinyol, M., et al. (2012). Epigenomic analysis detects widespread gene-body DNA hypomethylation in chronic lymphocytic leukemia. Nat Genet 44, 1236–1242.

Lister, R., Pelizzola, M., Dowen, R.H., Hawkins, R.D., Hon, G., Tonti-Filippini, J., Nery, J.R., Lee, L., Ye, Z., Ngo, Q.M., et al. (2009). Human DNA methylomes at base resolution show widespread epigenomic differences. Nature 462, 315–322.

Maianski, N.A., Maianski, A.N., Kuijpers, T.W., and Roos, D. (2004). Apoptosis of neutrophils. Acta Haematol 111, 56–66.

Maianski, N.A., Mul, F.P., van Buul, J.D., Roos, D., and Kuijpers, T.W. (2002). Granulocyte colony-stimulating factor inhibits the mitochondria-dependent activation of caspase-3 in neutrophils. Blood 99, 672–679.

Mantovani, A., Cassatella, M.A., Costantini, C., and Jaillon, S. (2011). Neutrophils in the activation and regulation of innate and adaptive immunity. Nat Rev Immunol 11, 519–531

Marshak-Rothstein, A. (2006). Toll-like receptors in systemic autoimmune disease. Nat Rev Immunol 6, 823–835.

Martens, J.H., and Stunnenberg, H.G. (2013). BLUEPRINT: mapping human blood cell epigenomes. Haematologica 98, 1487–1489.

Martinod, K., Demers, M., Fuchs, T.A., Wong, S.L., Brill, A., Gallant, M., Hu, J., Wang, Y., and Wagner, D.D. (2013). Neutrophil histone modification by peptidylarginine deiminase 4 is critical for deep vein thrombosis in mice. Proc Natl Acad Sci U S A 110, 8674–8679

McLean, C.Y., Bristor, D., Hiller, M., Clarke, S.L., Schaar, B.T., Lowe, C.B., Wenger, A.M., and Bejerano, G. (2010). GREAT improves functional interpretation of cis-regulatory regions. Nat Biotechnol 28, 495–501.

Neph, S., Stergachis, A.B., Reynolds, A., Sandstrom, R., Borenstein, E., and Stamatoyannopoulos, J.A. (2012). Circuitry and dynamics of human transcription factor regulatory networks. Cell 150, 1274–1286.

Nie Z, Hu G, Wei G, Cui K, Yamane A, Resch W, Wang R, Green DR, Tessarollo L, Casellas R, Zhao K, Levens D (2012). c-Myc is a universal amplifier of expressed genes in lymphocytes and embryonic stem cells. Cell 151.1: 68–79

Orecchia, A., Scarponi, C., Di Felice, F., Cesarini, E., Avitabile, S., Mai, A., Mauro, M.L., Sirri, V., Zambruno, G., Albanesi, C., et al. (2011). Sirtinol treatment reduces inflammation in human dermal microvascular endothelial cells. PLoS One 6, e24307.

Ostuni R, Natoli G, Cassatella MA, Tamassia N. Epigenetic regulation of neutrophil development and function. Semin Immunol. 2016 Apr;28(2):83–93.

Papayannopoulos, V., Metzler, K.D., Hakkim, A., and Zychlinsky, A. (2010). Neutrophil elastase and myeloperoxidase regulate the formation of neutrophil extracellular traps. J Cell Biol 191, 677–691.

Pfaff, D., Heroult, M., Riedel, M., Reiss, Y., Kirmse, R., Ludwig, T., Korff, T., Hecker, M., and Augustin, H.G. (2008). Involvement of endothelial ephrin-B2 in adhesion and transmigration of EphB-receptor-expressing monocytes. J Cell Sci 121, 3842–3850.

Pham, T.H., Benner, C., Lichtinger, M., Schwarzfischer, L., Hu, Y., Andreesen, R., Chen, W., and Rehli, M. (2012). Dynamic epigenetic enhancer signatures reveal key transcription factors associated with monocytic differentiation states. Blood 119, e161–171.

Pham, T.H., Minderjahn, J., Schmidl, C., Hoffmeister, H., Schmidhofer, S., Chen, W., Langst, G., Benner, C., and Rehli, M. (2013). Mechanisms of in vivo binding site selection of the hematopoietic master transcription factor PU.1. Nucleic Acids Res 41, 6391–6402.

Randolph, G.J., Jakubzick, C., and Qu, C. (2008). Antigen presentation by monocytes and monocyte-derived cells. Curr Opin Immunol 20, 52–60.

Reiss, M., and Roos, D. (1978). Differences in oxygen metabolism of phagocytosing monocytes and neutrophils. J Clin Invest 61, 480–488.

Robinson, P.N., and Mundlos, S. (2010). The human phenotype ontology. Clin Genet 77, 525–534.

Ronnerblad, M., Andersson, R., Olofsson, T., Douagi, I., Karimi, M., Lehmann, S., Hoof, I., de Hoon, M., Itoh, M., Nagao-Sato, S., et al. (2014). Analysis of the DNA methylome and transcriptome in granulopoiesis reveals timed changes and dynamic enhancer methylation. Blood 123, e79–89.

Salpea, P., Russanova, V.R., Hirai, T.H., Sourlingas, T.G., Sekeri-Pataryas, K.E., Romero, R., Epstein, J., and Howard, B.H. (2012). Postnatal development-and age-related changes in DNA-methylation patterns in the human genome. Nucleic Acids Res 40, 6477–6494.

Satterlee, J.S., Schubeler, D., and Ng, H.H. (2010). Tackling the epigenome: challenges and opportunities for collaboration. Nat Biotechnol 28, 1039–1044.

Schmidl, C., Renner, K., Peter, K., Eder, R., Lassmann, T., Balwierz, P.J., Itoh, M., NagaoSato, S., Kawaji, H., Carninci, P., et al. (2014). Transcription and enhancer profiling in human monocyte subsets. Blood 123, e90–99.

Shen, H., Qiu, C., Li, J., Tian, Q., and Deng, H.W. (2013). Characterization of the DNA methylome and its interindividual variation in human peripheral blood monocytes. Epigenomics 5, 255–269.

Stadler, M.B., Murr, R., Burger, L., Ivanek, R., Lienert, F., Scholer, A., van Nimwegen, E., Wirbelauer, C., Oakeley, E.J., Gaidatzis, D., et al. (2011). DNA-binding factors shape the mouse methylome at distal regulatory regions. Nature 480, 490–495.

Stunnenberg HG; International Human Epigenome Consortium, Hirst M. The International Human Epigenome Consortium: A Blueprint for Scientific Collaboration and Discovery. Cell. 2016 Nov 17;167(5):1145–1149.

Tilgner, H., Knowles, D.G., Johnson, R., Davis, C.A., Chakrabortty, S., Djebali, S., Curado, J., Snyder, M., Gingeras, T.R., and Guigo, R. (2012). Deep sequencing of subcellular RNA fractions shows splicing to be predominantly co-transcriptional in the human genome but inefficient for lncRNAs. Genome Res 22, 1616–1625.

Wang, G.G., Cai, L., Pasillas, M.P., and Kamps, M.P. (2007). NUP98-NSD1 links H3K36 methylation to Hox-A gene activation and leukaemogenesis. Nat Cell Biol 9, 804–812.

Wong, J.J., Ritchie, W., Ebner, O.A., Selbach, M., Wong, J.W., Huang, Y., Gao, D., Pinello, N., Gonzalez, M., Baidya, K., et al. (2013). Orchestrated intron retention regulates normal granulocyte differentiation. Cell 154, 583–595.

Zhou, L., Braat, H., Faber, K.N., Dijkstra, G., and Peppelenbosch, M.P. (2009). Monocytes and their pathophysiological role in Crohn’s disease. Cell Mol Life Sci 66, 192–202.

Ziller, M.J., Muller, F., Liao, J., Zhang, Y., Gu, H., Bock, C., Boyle, P., Epstein, C.B., Bernstein, B.E., Lengauer, T., et al. (2011). Genomic distribution and inter-sample variation of non-CpG methylation across human cell types. PLoS Genet 7, e1002389.

Zula, J.A., Green, H.C., Ransohoff, R.M., Rudick, R.A., Stark, G.R., and van Boxel-Dezaire, A.H. (2011). The role of cell type-specific responses in IFN-beta therapy of multiple sclerosis. Proc Natl Acad Sci U S A 108, 19689–19694.

## Supplementary references

1. S. Anders, W. Huber, Genome Biol 11, R106 (2010).

2. P. Flicek et al., Nucleic Acids Res, (Dec 6, 2013).

3. J. Harrow et al., Genome Res 22, 1760 (Sep, 2012).

4. L. A. Hindorff et al., Arch Intern Med 169, 68 (Jan 12, 2009).

5. S. John et al., Curr Protoc Mol Biol Chapter 27, Unit 21 27 (Jul, 2013).

6. D. Karolchik et al., Nucleic Acids Res, (Nov 21, 2013).

7. M. T. Maurano et al., Science 337, 1190 (Sep 7, 2012).

8. M. T. Maurano et al., Science 337, 1190 (Sep 7, 2012).

9. S. B. Montgomery et al., Nature 464, 773 (Apr 1, 2010).

10. A. Mortazavi, B. A. Williams, K. McCue, L. Schaeffer, B. Wold, Nat Methods 5, 621 (Jul, 2008).

11. S. Neph et al. Cell 150, 1274 (Sep 14, 2012).

12. M. D. Robinson, D. J. McCarthy, G. K. Smyth, Bioinformatics 26, 139 (Jan 1, 2010).

13. R. E. Thurman et al., Nature 489, 75 (Sep 6, 2012).

14. S. J. van Heeringen, G. J. Veenstra, Bioinformatics 27, 270 (Jan 15, 2011).

15. Y. Assenov et al., (2013) http://rnbeads.mpi-inf.mpg.de

16. S. Heinz et al., Mol Cell 38(4), 576–589 (May 28, 2010).

